# An immortalised mesenchymal stem cell line maintains mechano-responsive behaviour and can be used as a reporter of substrate stiffness

**DOI:** 10.1101/269225

**Authors:** Asier Galarza Torre, Joshua E. Shaw, Amber Wood, Hamish T. J. Gilbert, Oana Dobre, Paul Genever, Keith Brennan, Stephen M. Richardson, Joe Swift

## Abstract

The mechanical environment can influence cell behaviour, including changes to transcriptional and proteomic regulation, morphology and, in the case of stem cells, commitment to lineage. However, current tools for characterizing substrates’ mechanical properties, such as atomic force microscopy (AFM), often do not fully recapitulate the length and time scales over which cells ‘feel’ substrates. Here, we show that an immortalised, clonal line of human mesenchymal stem cells (MSCs) maintained responsiveness to substrate mechanics observed in primary cells, and could be used as a reporter of stiffness. MSCs were cultured on soft and stiff polyacrylamide hydrogels. In both primary and immortalised MSCs, stiffer substrates promoted increased cell spreading, expression of lamin-A/C and translocation of mechano-sensitive proteins YAP1 and MKL1 to the nucleus. Stiffness was also found to regulate transcriptional markers of lineage. A GFP-YAP / RFP-H2B reporter construct was designed and virally delivered to the immortalised MSCs for *in situ* detection of substrate stiffness. MSCs with stable expression of the reporter showed GFP-YAP to be increasingly co-localized with nuclear RFP-H2B on stiffer substrates, enabling development of a cellular reporter of substrate stiffness. This will facilitate mechanical characterisation of new materials developed for applications in tissue engineering and regenerative medicine.

## Introduction

Mechanical homeostasis is a fundamental property inherent to all tissues of the adult body. Establishment of the right stiffness for each tissue and stage in development is vital for the correct function of various organs^1^: bones, for example, must be stiff, while skin must be reversibly deformable. In order to maintain homeostasis in surrounding tissue, cells have mechanisms that allow them to ‘feel’ the mechanical properties of the extracellular matrix (ECM) and respond accordingly. Cells process physical stimuli through a set of mechanotransduction pathways, such as mechanically-regulated ion channels^2^ or focal adhesion (FA) complexes that assemble at the plasma membrane where cells pull on the surrounding ECM^3^. Mechanical signals are propagated within cells through pathways such as RhoA (Ras homolog gene family, member A) and ROCK (Rho-associated protein kinase) signalling^4^, and through regulation of transcription factors (TFs). Stiff substrates cause TFs such as YAP1 (yes-associated protein 1)^5^ and MKL1 (myocardin-like protein 1, also known as MRTF-A or MAL)^6^ to translocate to the nucleus, thus modulating their activity. Mechanical signals may also be transmitted through cells by a system of interlinked structural proteins that connect the ECM through FAs to the cytoskeleton, and then to the nucleus through the linker of nucleo- and cyto-skeleton (LINC) complex^7^. Mechanical inputs can therefore be passed from substrate to nucleus where they can affect chromatin conformation and thus influence how genes are regulated^8^.

A broad range of cellular processes have been shown to be influenced by mechanical inputs. These include the regulation of cell morphology^9^, such as the extent to which cells spread when adhering to a twodimensional substrate, and the amount of force that cells apply to deform their surroundings^10^. Changes to cell morphology and contractility require regulation of protein content within the cells, and this has been characterised in the cytoskeleton and the nuclear lamina^11^. Apoptosis pathways and the rate of proliferation are also influenced by substrate stiffness^12^, and cells such as fibroblasts have been shown to migrate along gradients of increasing stiffness, a process called durotaxis^13^. Mesenchymal stem cells (MSCs) have been used as a model system to examine a number of mechanotransduction processes^4,5,11,14^, with sensitivity to mechanical stimulation noted in even seminal characterisations^15^. MSCs are multipotent cells with lineage potential that can be influenced by substrate mechanics^16^: culture on soft substrates favours adipogenesis, while stiff substrates favour osteogenesis.

The multipotent nature of MSCs combined with a capacity to modulate immune responses^17^ have led to investigations of their suitability for regenerative medicine, and the possibility of replacing damaged tissues with engineered scaffolds repopulated with stem cells^18,19^. James *et al.* have reported the generation of immortalised MSCs through overexpression of telomerase reverse transcriptase (TERT)^20^. Clonal lines were selected for expansion that showed exponential growth, matched the expected cell-surface marker profile for MSCs and presented no evidence of tumorigenicity in immuno-compromised mice. Endogenous populations of MSCs are typically heterogeneous, but the selected clonal lines exhibited specific characteristics of MSC behaviour. Some of the clones exhibited immuno-modulatory behaviour, while the clonal line “Y201” maintained potential for chemical induction of adipogenic, chondrogenic and osteogenic lineages. Here we compared the responses of primary and Y201 MSCs to culture on soft and stiff collagen-I coated polyacrylamide (PA) hydrogels to determine the extent to which mechanosensitive behaviour was maintained in the immortalised line. We found that the Y201 MSCs preserved the mechanosensitive features of primary cells: morphological responses such as cell spreading on stiffer substrates, the ability to remodel the nucleoskeleton, translocate TFs and regulate genes associated with adipogenesis and osteogenesis.

Recent efforts in the field of tissue engineering have sought to develop substrates and scaffolds on which cells can be cultured for use in therapy or to produce synthetic tissues that can ultimately be implanted into patients. Substrates have been generated using diverse chemistries, enabling properties such as tuneable stiffness^21–24^ or the incorporation of ECM molecules^25^. Scaffolds have been constructed using a variety of technologies, from bio-inspired, synthetic polymers^26^ to human-derived, decellularised tissues^27^. The behaviour of cells seeded onto such materials is influenced by the combined chemical and mechanical characteristics of the microenvironment^28^. It is therefore often desirable to characterise the mechanical properties of culture substrates and scaffolds. Atomic force microscopy (AFM), nanoindentation and rheology offer means of measuring material parameters, such as the elasticity (Young’s modulus, *E*), but are in many ways limited. Cell behaviour may be influenced by deformations over specific length or time scales, by complex viscoelastic material properties, by heterogeneity, or by dimensionality and topologies that would be difficult to examine with a mechanical probe (such as an AFM tip) or bulk measurement method (such as rheology). The best reporter of how cells respond to a particular mechanical environment - and correspondingly to direct how materials can be developed to control cell behaviour - may therefore be to use a probe technology based on the cells themselves. With this aim, having established the mechanosensitivity of the Y201 MSCs, we used them as a platform to develop a fluorescent stiffness probe. Here we demonstrate the use of immortalised MSCs stably expressing a red/green fluorescent reporter construct that can be analysed by microscopy to differentiate soft and stiff substrates.

## Results

### Immortalised MSCs maintain a morphological response to substrate stiffness

In order to determine whether an immortalised MSC cell line, previously reported to show the multipotent capacity of primary cells^20^, also maintained sensitivity to mechanical inputs, the response of the cells to culture on soft or stiff substrates was characterised. Immortalised MSCs, along with a comparison set of primary cells from four human donors, were cultured on soft (2 kPa) or stiff (25 kPa) collagen-I coated PA hydrogels for three days, before being fixed and stained with DAPI and phalloidin (Fig. 1a). Cells on stiffer substrates appeared larger with more pronounced and ordered stress fibres. Quantitative analysis of cell morphology showed that the mean spread area of primary MSCs was increased 1.4-fold (*p* = 0.007; *n* = 4 donors) on stiff versus soft hydrogels; the spread area of the immortalised line was increased 1.5-fold under the same conditions (*p* < 0.0001; Fig. 1b). Cell aspect ratio (defined as the ratio of long to short side lengths of the smallest rectangle that can enclose the perimeter of a cell) was not significantly affected by substrate stiffness in either primary or immortalised cells (Fig. 1c). Cell circularity (proportional to the ratio between the area and the square of the perimeter of a cell) was significantly lower on the stiff substrate for primary cells (*p* = 0.004; *n* = 4 donors), but not significantly altered in immortalised cells (Fig. 1d). Analysis of nuclear morphology in the same samples yielded a parallel set of characteristics. The spread areas of the nuclei of primary cells were increased 1.1-fold in both primary and immortalised cells, although this effect was only significant in the immortalised line (*p* < 0.0001), perhaps reflecting the greater variation amongst cells from primary donors (Fig. 1e). Nuclear aspect ratio was significantly increased on stiff substrates in both primary and immortalised MSCs (*p* = 0.003 and *p* < 0.0001, respectively; Fig. 1f). Nuclear circularity was not affected by stiffness in primary cells, although it was fractionally reduced on stiff substrates in the immortalised line (p < 0.0001; Fig. 1g). These results established that immortalised MSCs broadly replicated the characteristic morphological responses of primary cells when subjected to modulated substrate mechanics.

**Figure 1.**
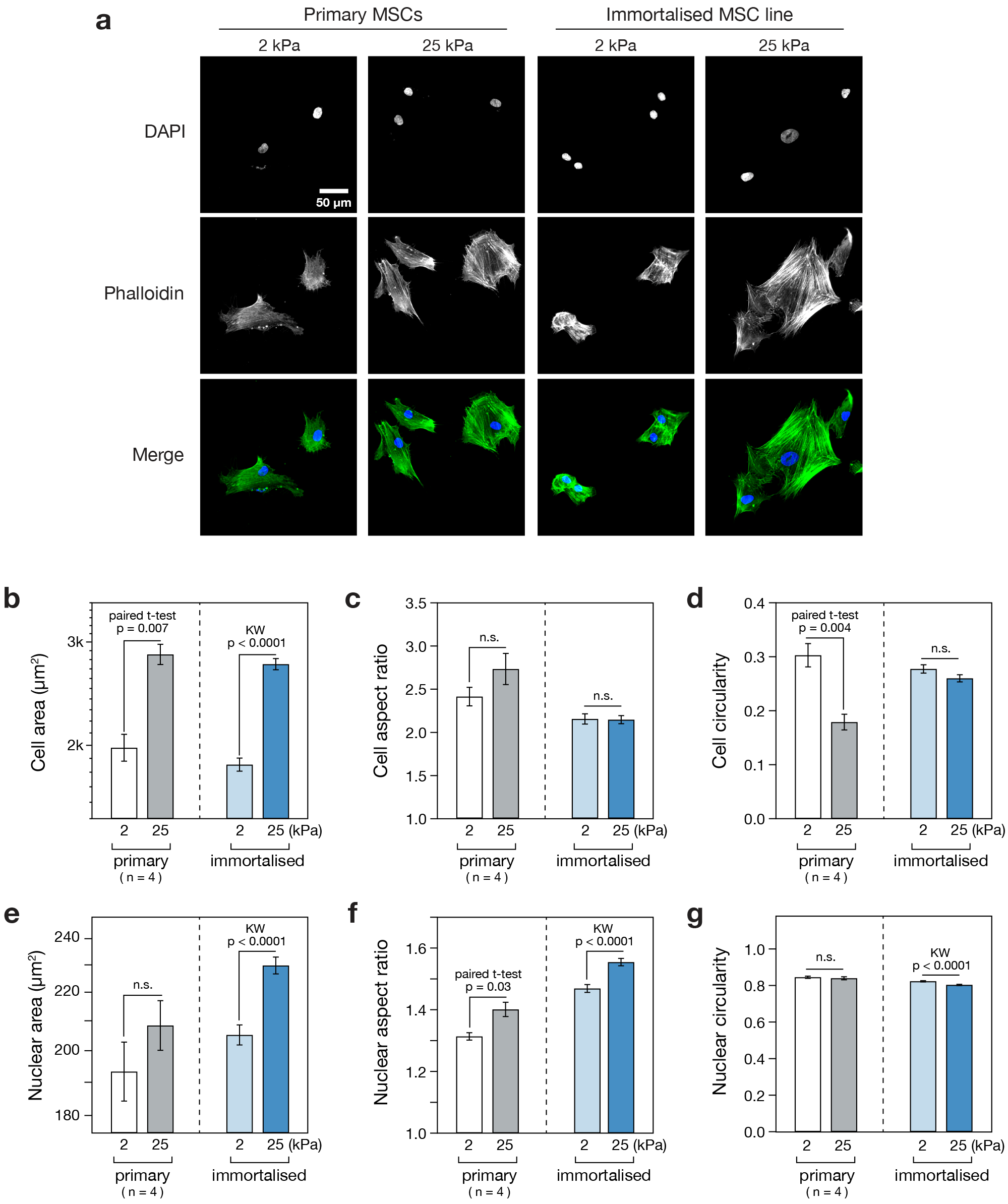
Cellular and nuclear morphology are stiffness-responsive in primary and immortalised MSCs. (a) MSC morphology was assessed using phalloidin and DAPI staining following 3 days culture on soft (2 kPa) or stiff (25 kPa) collagen-I coated PA hydrogels. (b) The spread areas of both primary and immortalised MSCs were significantly greater on stiff substrates than on soft. (c) Cell aspect ratios were not significantly affected by stiffness. (d) Cell circularity was significantly decreased in primary MSCs on stiff substrates; this effect was not observed in immortalised cells. (e) Nuclear area was increased on stiff substrates, although this effect was only significant in immortalised MSCs. (f) Nuclear aspect ratio was significantly increased on stiff substrates in both primary and immortalised MSCs. (g) Nuclear circularity was not significantly affected by increased stiffness in primary MSCs, but was lowered in immortalised MSCs (p-values from paired t-tests and Kruskal-Wallis (KW) tests as indicated; n.s. = not significant; *n* = 4 primary donors; a minimum of 38 cells were analysed per condition for each primary donor; a minimum of 363 immortalised cells were analysed per condition).

### The nuclear lamina of primary and immortalised MSCs is stiffness-responsive

The composition of the nuclear lamina has been shown to be affected by substrate stiffness^11^. Primary and immortalised MSCs were cultured on soft (2 kPa) and stiff (25 kPa) collagen-I coated PA hydrogels for three days before being fixed and imaged with immuno-staining against lamins A/C and B1 (Fig. 2a). In order to compensate for changes in nuclear spread area, the ratio of lamin A/C to B1 stain intensity within the nucleus was calculated. Lamin B1 was used for normalisation as it has been reported to be relatively unresponsive to substrate stiffness^11^. Analysis of MSCs from four primary donors, in comparison to the immortalised line, showed that the response to stiffness was maintained: the LMNA:B1 ratio was significantly greater on the stiffer substrate (*p* < 0.0001; Fig. 2b). The mean LMNA:B1 ratio was increased 2.2-fold on stiff versus soft substrates in the primary MSCs (*p* = 0.02; *n* = 4 donors); in the immortalised MSCs, the increase was 2-fold (*p* < 0.0001; Fig. 2c). This result suggests that the mechanosensitivity of cell adhesions, and consequent cell spreading with increased substrate stiffness, was propagated to the nucleus in primary MSCs, consistent with earlier reports^11^, and that the mechanism was preserved in the immortalised cell line.

**Figure 2.**
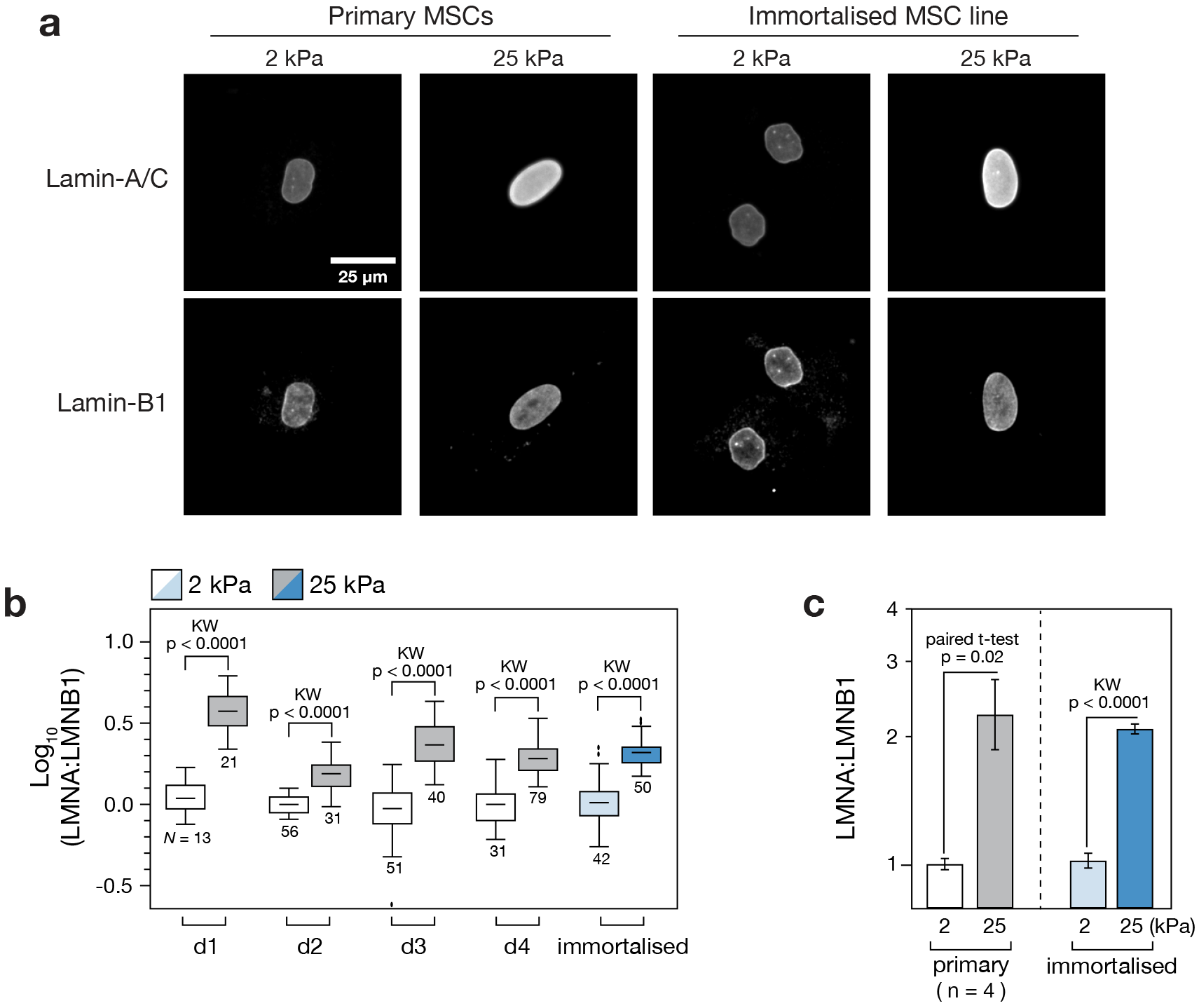
Composition of the nuclear lamina is stiffness-responsive in primary and immortalised MSCs. (a) Immunofluorescence imaging was used to quantify lamins -A/C and -B1 (LMNA and LMNB1) in primary and immortalised MSCs cultured on soft (2 kPa) or stiff (25 kPa) collagen-I coated PA hydrogels. Lamin-B1 expression appeared constant in all conditions, consistent with earlier reports^11^. (b) Quantification of lamin A/C to B1 ratio (LMNA:LMNB1) from immunofluorescence images showed that lamin-A/C was consistently upregulated on stiff substrates, relative to lamin-B1, in each of four primary donors and the immortalised MSCs (N indicates number of cells analysed per condition). (c) LMNA:LMNB1 was significantly increased on stiff substrates (p-values from paired t-tests and Kruskal-Wallis (KW) tests as indicated; *n* = 4 primary donors).

### Immortalised MSCs maintain the mechano-sensitive translocation of transcription factors YAP1 and MKL1

Primary and immortalised MSCs cultured on soft (2 kPa) and stiff (25 kPa) hydrogels for three days were also examined with immunofluorescence microscopy to determine the subcellular localisation of TFs YAP1 and MKL1, previously reported to translocate to the nucleus on stiff substrates^5,6^. YAP1 was found to be increasingly localised to the nucleus on the stiffer hydrogel in both primary and immortalised MSCs (Fig. 3a). Quantification of the ratio of nuclear to cytoplasmic YAP1 (determined by using DAPI and phalloidin channels to define cell and nuclear boundaries) showed donor-to-donor variation across four samples (Fig. 3b). Nonetheless, YAP1 was significantly more localised to the nucleus in three of the four samples (*p* < 0.0001). The mean nuclear:cytoplasmic ratio of YAP1 increased 1.7-fold on stiff versus soft substrates in the primary MSCs (not significant, *p* = 0.08; *n* = 4 donors) while the increase was 1.3-fold (*p* < 0.0001) in the immortalised cells (Fig. 3c). Mean total YAP1 (the sum of nuclear and cytoplasmic protein) was significantly downregulated on stiff substrates in the primary MSCs (*p* = 0.02), but was not affected in the immortalised cells (Fig. 3d). Examination of samples prepared in parallel showed the distribution of MKL1 to be similarly affected by substrate mechanics (Fig. 3e). The nuclear:cytoplasmic ratio of MKL1 was significantly higher on stiff than soft substrates in three of the four primary donor samples (*p* < 0.0009; Fig. 3f). Interestingly, the cells from donor 2 (d2) that showed no mechano-responsiveness in YAP1 cellular location also showed no significant regulation of MKL1. This reflects the variability among primary MSCs from different donors, and suggests the potential experimental utility of an immortalised line that reproduces canonical mechanosensitive features. The mean nuclear:cytoplasmic ratio of MKL1 was increased 1.5-fold in the primary cells (*p* = 0.04; *n* = 4 donors) and 1.3-fold in the immortalised line (*p* < 0.0001; Fig. 3g). In contrast to YAP1, total MKL1 was highly responsive to substrate stiffness, increasing 2.7-fold on stiff substrates in the primary cells (*p* = 0.03; *n* = 4 donors) and 5.5-fold in the immortalised cells (*p* < 0.0001; Fig. 3h). Both primary and immortalised MSCs thus exhibited substrate-directed regulation of TF subcellular localization.

**Figure 3.**
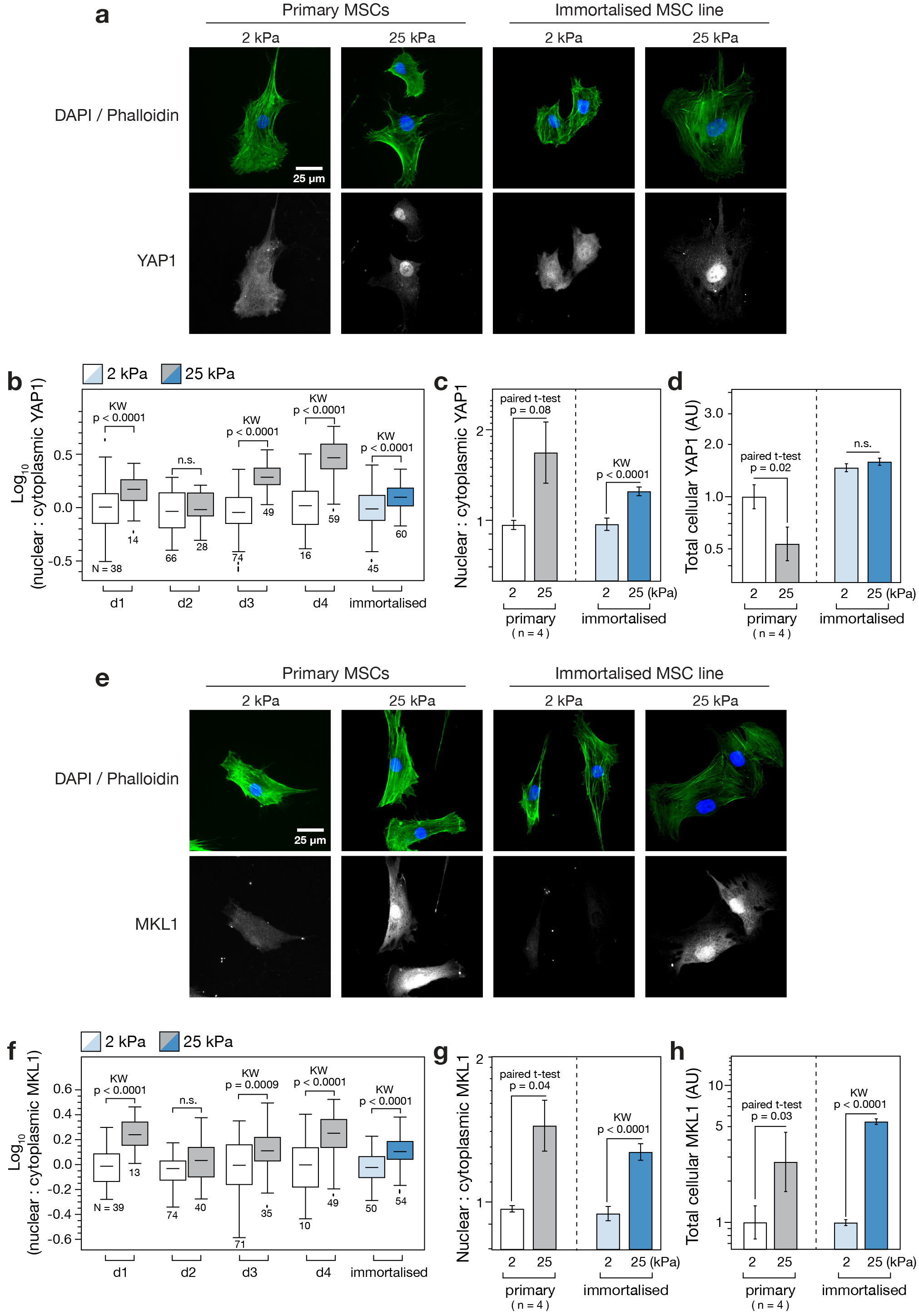
Transcription factors YAP1 and MKL1 respond to stiffness in primary and immortalised MSCs. (a) Immunofluorescence imaging was used to examine the location of yes-associated protein 1 (YAP1) in primary and immortalised MSCs cultured on soft (2 kPa) or stiff (25 kPa) collagen-I coated PA hydrogels. The nucleus and extent of the cytoplasm were identified by DAPI and phalloidin staining respectively. (b) YAP1 became increasingly localised in the nucleus on stiff substrates in MSCs from three of four primary donors, and in immortalised MSCs (N indicates number of cells analysed per condition). (c) Relative nuclear localisation of YAP1 was significantly increased in immortalised MSCs on stiff substrates. (d) The total amount of YAP1 (integrated signal from the whole cell) was significantly lower on stiff substrates in primary cells, but unchanged in immortalised cells. (e) Cellular location of myocardin-like protein 1 (MKL1, also known as MRTF-A or MAL) was imaged by immunofluorescence in primary and immortalised MSCs on soft and stiff substrates. (f) MKL1 was increasingly localised in the nucleus on stiff substrates in MSCs from three of four primary donors, and in immortalised MSCs (*N* indicates number of cells analysed per condition). (g) MKL1 was significantly more localised to the nucleus on stiff substrates in primary and immortalised cells (h) Total levels of MKL1 were also highly dependent on substrate stiffness in both primary and immortalised cells: in both cases, MKL1 was significantly higher on stiff substrates (*p*- values from paired t-tests and Kruskal-Wallis (KW) tests as indicated; n.s. = not significant; *n* = 4 primary donors).

### Substrate stiffness modulates lineage markers CEBPA and RUNX2 in immortalised MSCs

Primary MSCs cultured for three weeks on tissue culture plastic (TCP) in media supplemented with an adipo-induction cocktail formed lipid droplets that could be imaged by Oil Red O staining. Consistent with previous reports^20^, the immortalised MSCs also formed lipid droplets under the same conditions (Fig. 4a). The influence of substrate stiffness alone was sufficient to alter transcript levels of adipogenic marker *CEBPA* (CCAAT/enhancer-binding protein alpha) in immortalised MSCs: *CEBPA* was 2.4-fold higher in cells cultured on soft (2 kPa) versus stiff (25 kPa) collagen-I coated PA hydrogels in standard media for three weeks (*p* = 0.0006; Fig. 4b). Regulator of adipocyte differentiation *PPARG* (Peroxisome proliferator-activated receptor gamma) was also 1.3-fold higher on soft than stiff substrates, but this effect was not significant. Modulation of substrate stiffness did not significantly affect levels of *CEBPA* or *PPARG* transcripts in immortalised cells treated with adipo-induction media (Fig. 4c). Culture of primary MSCs on TCP for three weeks in media supplemented with an osteo-induction cocktail showed positive staining for alkaline phosphatase (ALP) activity. As reported previously^20^, the immortalised line also showed ALP activity under the same conditions (Fig. 4d). The influence of substrate stiffness was again found to be sufficient to affect genes indicative of lineage in the absence of chemical induction: culture of immortalised MSCs on stiff substrates caused a 4-fold increase in *RUNX2* (runt-related transcription factor 2) relative to culture on soft (*p* = 0.03; Fig. 4e). *BGLAP* (osteocalcin) was also increased, but the change was not significant. *RUNX2* transcript was increased 16-fold on stiff versus soft in immortalised MSCs cultured with osteo-induction media (*p* = 0.02), suggesting a synergistic relationship between chemical and mechanical stimuli (Fig. 4f). These results reproduce earlier observations of differentiation potential in the clonal, immortalised Y201 MSC line and furthermore show that genes associated with adipogenesis and osteogenesis are mechanically regulated.

**Figure 4.**
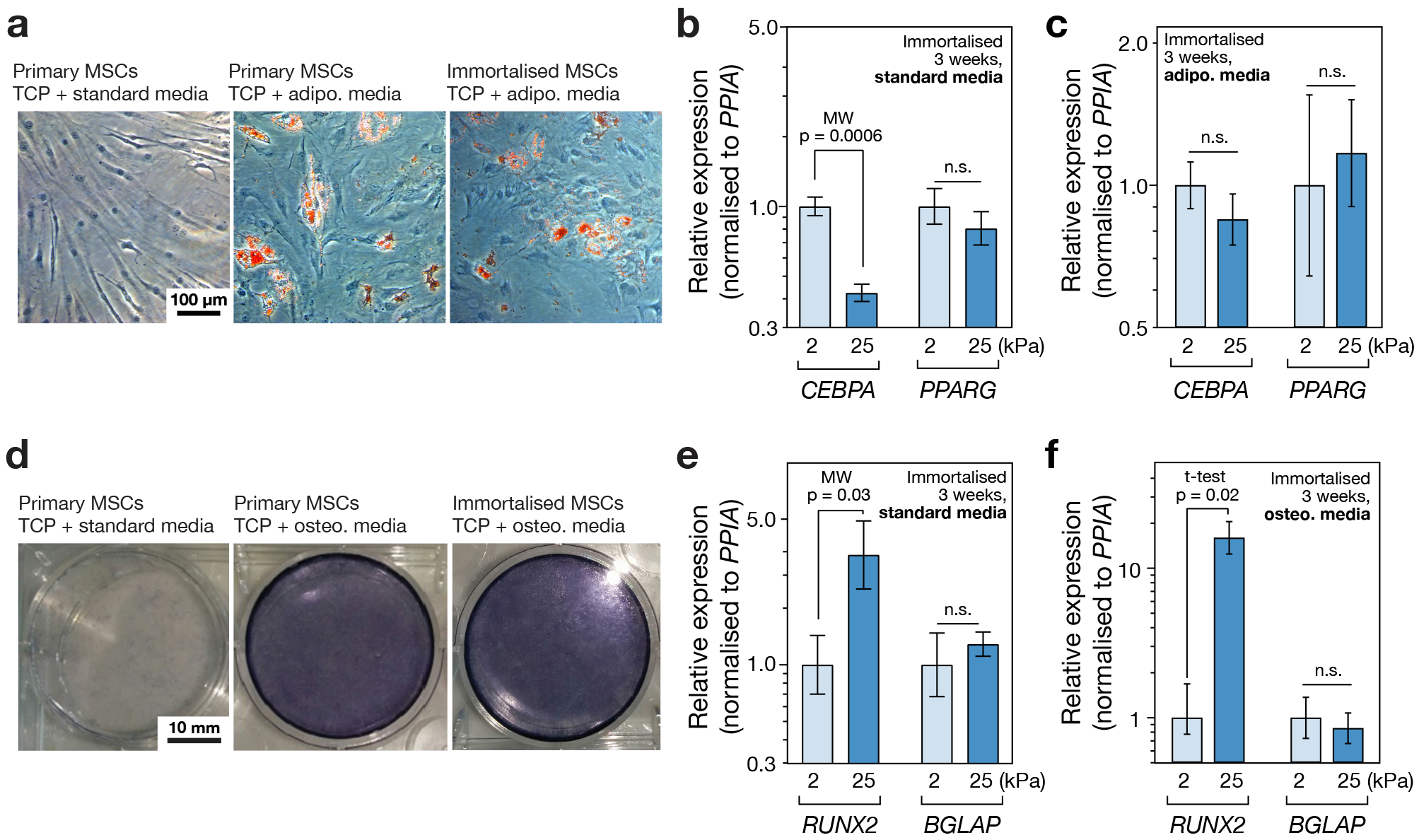
(a) Images of MSCs cultured on tissue culture plastic (TCP) for three weeks in the presence of adipogenic induction media, with Oil Red O staining (in red). Both primary and immortalised MSCs showed positive staining of lipid droplets, indicating adipogenic potential and confirming observations reported previously^20^. (b) Culture on soft (2 kPa) collagen-I coated PA hydrogels significantly increased the adipogenic marker *CEBPA*, relative to culture on stiff (25 kPa) hydrogels, measured by RT-qPCR in immortalised MSCs after three weeks in standard media. A second adipogenic marker *PPARG* was also higher on soft substrates, although this effect was not significant. (c) Combining substrate stiffness cues with adipogenic induction media did not significantly change levels of *CEBPA* or *PPARG*. (d) MSCs cultured on TCP for three weeks in the presence of osteogenic induction media and stained for alkaline phosphatase activity (ALP, in blue). Both primary and immortalised MSCs showed positive ALP staining, indicating an osteogenic potential in agreement with earlier reports^20^. (e) Culture on stiff PA hydrogels significantly increased the osteogenic marker *RUNX2*, relative to culture on soft, measured in immortalised MSCs after three weeks in standard media. Osteogenic marker *BGLAP* was also higher on stiff substrates, although this effect was not significant. (f) Stiff substrate significantly amplified the effect of chemical osteogenic induction on *RUNX2* in immortalised MSCs; *BGLAP* levels were not significantly altered. A synergistic interaction between mechanical and chemical inputs in influencing lineage potential has previously been reported in primary MSCs^11^ (p-values from Mann-Whitney (MW) and un-paired t-tests as indicated; n.s. = not significant).

### A stably expressed reporter construct responds to substrate stiffness

The immortalised MSC line was transformed to stably express histone H2B labelled with red fluorescent protein (RFP-H2B) and either wildtype YAP1 tagged with green fluorescent protein (GFP-YAP(wt)) or a tagged YAP1 with four point mutations to serine residues (GFP-YAP(4SA); See supplemental Figure 1 for plasmid maps). The construct with the mutant YAP1 was considered here to investigate whether point mutations affecting protein turnover could produce a more stable reporter system. The reporter cells were cultured on soft (2 kPa) or stiff (25 kPa) substrates for three days under standard conditions, fixed and imaged (Fig. 5a). The RFP-H2B was localised to the nuclei of the MSCs; the GFP-YAP constructs appeared throughout the cell, but were observed with increasing intensity in the nuclei of MSCs on stiff substrates, particularly in the case of the wildtype YAP construct. The mechanosensitivity of cell spreading was maintained in MSCs with GFP-YAP(wt) and GFP-YAP(4SA) constructs (*p* = 0.0002 and 0.03, respectively; Fig. 5b), reflecting the observations made of primary and non-transformed cells (Fig. 1). Nuclear area also showed sensitivity to the stiffness of the substrate in the GFP-YAP(wt) MSCs (Fig. 5c). Quantification of the nuclear:cytoplasmic ratio of GFP-YAP(wt) showed the construct to be 1.5-fold more intensely localised to the nucleus in MSCs cultured on stiff (25 kPa) than on soft (2 kPa) hydrogels (*p* < 0.0001; Fig. 5d). The GFP-YAP(4SA) construct also maintained significant mechanosensitivity (*p* < 0.0001), although the fold change in nuclear:cytoplasmic ratio was decreased. The integrated, whole-cell intensity of GFP-YAP(wt) was significantly lower on stiff substrates (*p* = 0.03), but levels of the GFP-YAP(4SA) construct showed no significant sensitivity to stiffness (Fig. 5e).

**Figure 5.**
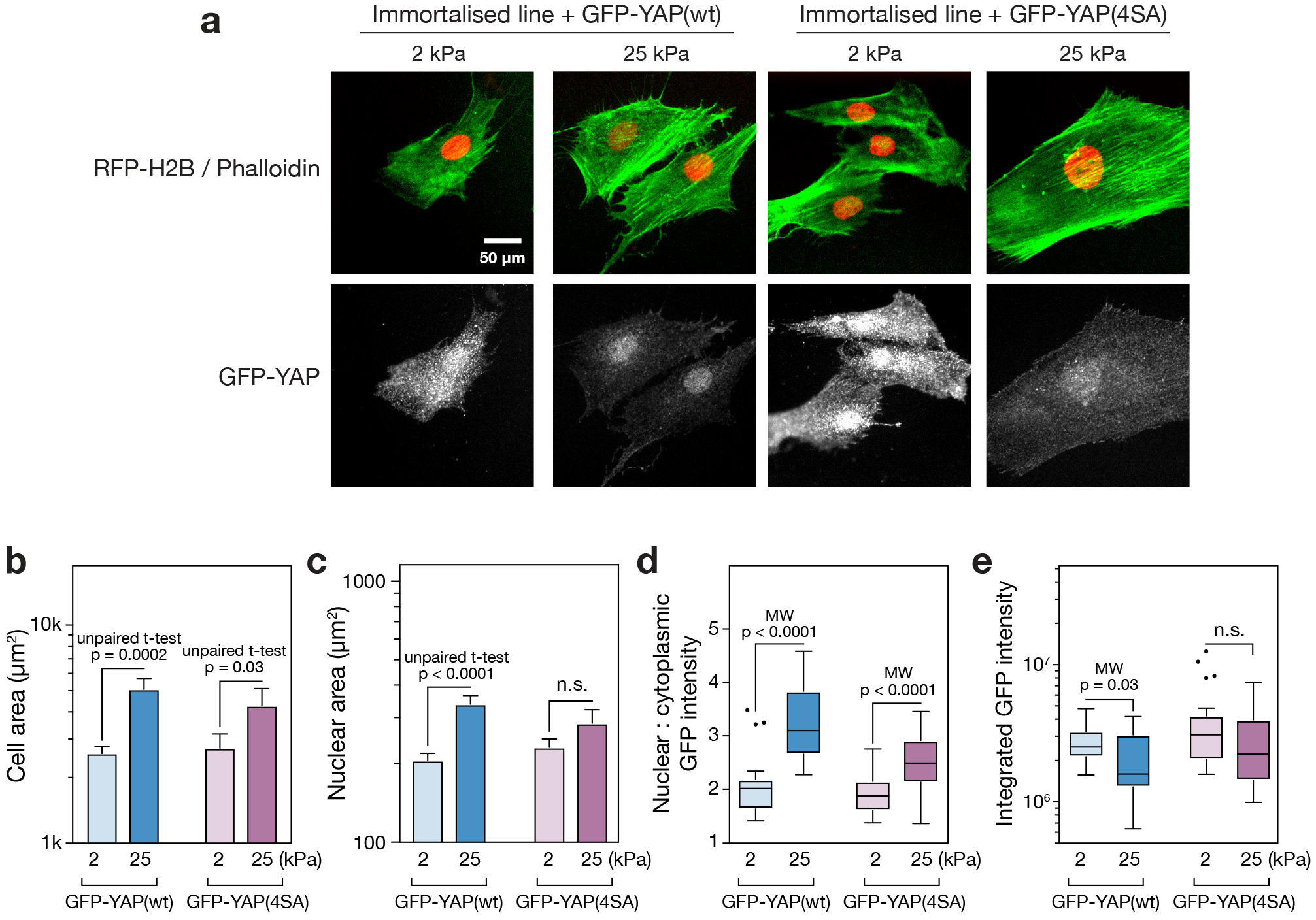
Application of a GFP-YAP/RFP-H2B fluorescent construct to report on substrate stiffness. (a) Immortalised MSCs were virally transformed to express RFP-H2B and either GFP-YAP(wt) (wild type) or GFP-YAP(4SA) (YAP1 modified with mutations to four serine residues^43^). Cells were cultured on soft (2 kPa) or stiff (25 kPa) hydrogels for three days, fixed and stained with phalloidin and anti-GFP. (b) Analysis of cell areas of the transformed immortalised MSCs showed that responsiveness to substrate mechanics was maintained: spread area was significantly greater on stiff (25 kPa) versus soft (2 kPa) hydrogels with both the GFP-YAP(wt) and GFP-YAP(4SA) construct. (c) Nuclear spread area of the GFP-YAP(wt) transformed cells was significantly greater on stiffer hydrogels; nuclear spread of GFP-YAP(4SA) was also greater, although not significant. (d) Image analysis showed that nuclear localisation of the YAP reporter constructs was higher on stiff (25 kPa) than soft (2 kPa) substrates. Translocation was significant for both GFP-YAP(wt) and GFP-YAP(4SA) constructs, although the magnitude of the effect was greater with the wildtype construct. (e) Total GFP-YAP(wt) was significantly lower on stiff substrates. Total GFP-YAP(4SA) was greater than in the wildtype construct, although not significantly so, and responsiveness to substrate stiffness was lost. (p-values from Mann-Whitney (MW) and un-paired t-tests as indicated; n.s. = not significant; outliers in box-whisker plots were identified by the Tukey method; a minimum of 17 cells were analysed under each condition).

## Discussion

### An immortalised MSC line maintains mechanosensitive characteristics of primary cells

Since the earliest characterisations, MSCs have been reported to be responsive to mechanical input^15^, and have subsequently been used as a model system to study a wide range of mechanosensitive phenomena^4,5,11,14^. The immortalised MSCs (clonal line Y201, initially isolated from human bone marrow) generated by James *et al.* have been shown to maintain the adipogenic, chondrogenic and osteogenic potential observed in primary MSCs when treated with standard, differentiation-inducing chemical cocktails^20^. However, the extent to which immortalised MSCs preserved their responsiveness to substrate mechanics - and, by extension, could be used as a platform to develop mechanosensing tools - had not been established. Here, primary and immortalised cells were cultured on soft and stiff collagen-I coated PA hydrogels and aspects of recognised mechanosensing pathways compared. Hydrogel stiffnesses were chosen to be representative of the bone marrow microenvironment: fat and marrow tissues are soft while precalcified bone is comparatively stiff, modelled by 2 and 25 kPa gels, respectively^11,29^.

The immortalised MSCs reproduced the characteristic morphological responses to substrate stiffness observed in primary MSCs (Fig. 1). Adherent cells pull on and probe the surrounding microenvironment^30^ and signalling pathways activated in FA complexes^1^ drive cell spreading through reorganisation of the actin cytoskeleton^31^. Cell morphology has also been shown to be an effective predictor of cell fate^32^. Primary and immortalised MSCs were found to spread more on the stiffer substrate (Fig. 1b), although cellular aspect ratio and circularity were more sensitive to substrate in primary cells (Figs. 1c and d). Cell elongation has previously been reported for MSCs cultured on substrates of around 10 kPa ^33^. The ECM, cytoskeleton and nucleoskeleton are considered to be part of a continuous, mechanically linked system^7,34^. Changes in cellular morphology were matched by coincident changes in nuclear morphology (Fig. 1e), suggesting that the LINC complex that tethers the cytoskeleton to the nucleoskeleton remained functional in the immortalised cells. The nucleoskeleton, and specifically an increase in the ratio of A-type to B-type lamins, has been shown to report on a more contractile cell phenotype consequent of a stiffer substrate^11^. The ratio of lamin A/C to B1 was found here to be significantly higher on stiff substrates in both primary and immortalised cells (Fig. 2). This result again suggested that mechano-transmission pathways remained functional in the immortalised line. This is important as a range of signalling pathways, such as serum response factor (SRF), have been shown to be downstream of lamin-A/C regulation^35^.

Translocation of TFs between cytoplasm and nucleoplasm is a common motif within mechanosensing pathways, allowing TF activity to be modulated and thus providing control over specific transcriptionally regulated programs, such as lineage selection^36,37^. Regulation of the subcellular localization of TFs YAP1 and MKL1 as a function of substrate stiffness was found to be maintained in the immortalised MSCs (Fig. 3). Stiffness-directed regulation of YAP1 has been shown to be necessary for mechanically influenced lineage determination in MSCs, with nuclear YAP1 driving osteogenesis^5^. The mechanical regulation of MKL1 and its influence on SRF have also been well characterised^38^, including through interactions with lamin-A/C and emerin^6^. MKL1 has also been shown to be important for PPARG mediated adipogenesis in pre-adipocyte cell lines^39^.

The differentiation potential of MSCs has been shown to be influenced by the mechanical properties of the microenvironment, with soft substrates favouring soft tissue specification, such as adipogenesis, and stiff substrates favouring osteogenesis^5,11,16^. Seminal characterisation of the Y201 immortalised MSC line showed that adipogenic lineage was induced by treatment with a standard adipo-induction cocktail, evidenced by positive Oil Red O staining and increased levels of *LPL* (lipoprotein lipase) and *PPARG* transcript after 14 and 21 days^20^. Here, chemically-induced adipogenesis was confirmed in immortalised MSCs cultured on TCP (Fig. 4a). Additionally, levels of the adipogenic reporter gene *CEBPA* were found to be increased in cells cultured on soft versus stiff substrates for 21 days in the absence of chemical stimulation (Fig. 4b). James *et al.* also reported chemically-induced osteogenesis in the Y201 line on TCP, coincident with increased levels of *ALP* and *RUNX2* transcripts^20^. In addition to reproducing positive ALP staining on TCP (Fig. 4d), *RUNX2* transcript was found here to be increased in the immortalised MSCs cultured on stiff versus soft hydrogels for 21 days in the absence of chemical stimulation (Fig. 4e). Furthermore, *RUNX2* was more substantially affected by substrate stiffness when the immortalised MSCs were cultured in the presence of the osteo-inducing chemical cocktail, suggesting a synergistic relationship between chemical and physical inputs (Fig. 4f). Taken together, these results show that the mechano-responsive nature of MSC behaviour was maintained in an immortalised clonal line, from initial responses in cell morphology driven by ECM-FA interactions, through mechano-transmission to the nuclear skeleton and modulation of TF activity, and to influence the gene regulation that can affect cell fate (Fig. 6a).

**Figure 6.**
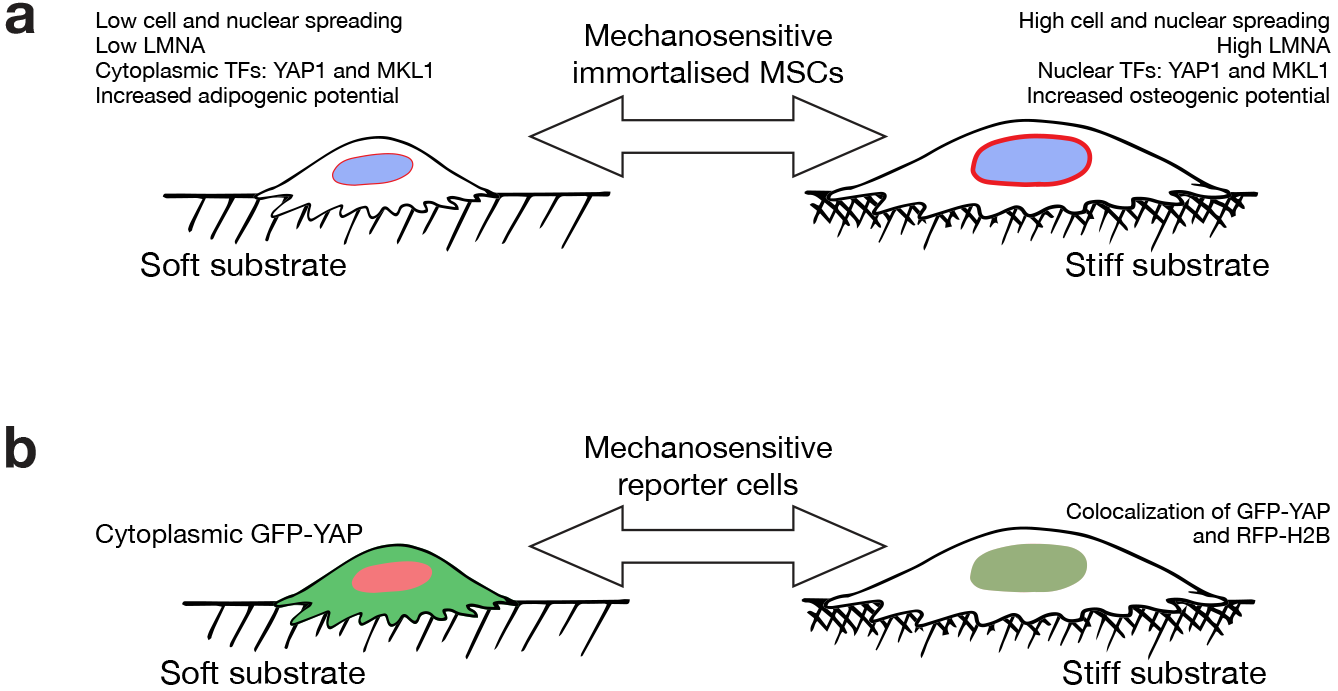
Mechanosensitive features of primary MSCs maintained in the immortalised MSC line. (a) Mechano-regulatory pathways observed in both primary and immortalised cells included: modulation of cell spreading and morphology, regulation of the nuclear lamina, translocation of transcription factors YAP1 and MKL1, and changes to differentiation potential. (b) The maintained sensitivity of YAP1 location to substrate stiffness enabled development of a reporter cell line: cytoplasmic GFP-YAP was indicative of a soft substrate, while co-localisation with nuclear RFP-H2B indicated a stiff substrate.

### Generation of a stable cell line with a fluorescent stiffness reporter

Here we showed that cells could be used as tools to report on substrate stiffness (Fig. 5). Using MSCs as a platform on which to base such a technology is rational given their well-characterised mechanoresponse and long-established use in regenerative medicine research^18,40^. However, we found the extent of mechanosensitivity to vary between cells from different primary donors (Figs. 2b, 3b and f) and variation has been described within heterogeneous populations derived from single donors^41^. Furthermore, a tool based on primary MSCs would have a limited lifespan as these cells typically show senescence over 8 - 12 passages^42^. The availability of an immortalised clonal MSC line (Y201)^20^ with maintained mechanosensing properties therefore represented an opportunity to develop a more reproducible measurement system. Two mechanosensitive TFs were identified as candidates on which to base a fluorescent stiffness probe. Both YAP1 and MKL1 maintained mechanosensitivity in the immortalised MSC line, translocating to the nucleus on stiff substrates (Figs. 3c and g). However, YAP1 was considered a better basis for a reporter construct as its mechanoresponse in the immortalised cells manifested only in its subcellular localisation, whereas both total MKL1 and its cellular compartment were affected by substrate stiffness (Figs. 3d and h).

The immortalised MSCs were transformed to express nuclear-localised, RFP-tagged histone H2B and wildtype YAP1 labelled with GFP. In addition, immortalised MSCs were transformed to express the RFP-H2B plus a GFP-tagged YAP1 with point mutations to four serine residues (S61A, S109A, S164A and S381G; “GFP-YAP(S4A)”). The intracellular location of both the GFP-YAP(wt) and GFP-YAP(S4A) probes was sensitive to substrate stiffness, being significantly more localised to the nucleus on a stiff (25 kPa) versus soft (2 kPa) hydrogel in both wildtype and mutant cases (Figs. 5a and d). YAP is phosphorylated by the serine/threonine protein kinase LATS (large tumour suppressor) at five serine residues as part of the Hippo signalling cascade. Phosphorylation at serine-127 - not mutated in the GFP-YAP(S4A) construct - is thought to regulate translocation between cytoplasm and the nucleus, while phosphorylation at serine-381 - mutated to a glycine in the GFP-YAP(S4A) construct - heads a cascade leading to proteasomal degradation of YAP^43^. It was therefore reasoned that the GFP-YAP(S4A) construct might retain the ability to report on stiffness because of its location within the cell, but be more stable and less liable to disturb other cellular functions. This was partially borne out by an increased stability of cytoplasmic GFP-YAP(S4A) that resulted in no significant change to total levels of the construct on soft and stiff substrates (Fig. 5e). However, the sensitivity and dynamic range of GFP-YAP(4SA) translocation was also reduced with respect to the wildtype construct (Fig. 5d). Previous reports have shown that overexpression of YAP can override the influence that substrate stiffness has over MSC lineage^5^. However, here the presence of the reporter constructs was found not to affect the initial mechanosensitive cell and nuclear spreading responses (Figs. 5b and c), found in the primary and untransformed immortalised MSCs (Fig. 1) and interpreted as being indicative of intact mechanosensing mechanisms at FAs. The GFP-YAP(wt) construct was thus found to translocate such that the nuclear:cytoplasmic ratio could be used to report on substrate stiffness (Fig. 6b).

In conclusion, we have shown that an immortalised, clonal line of human-derived MSCs maintained the mechanosensitive features of primary MSCs. Furthermore, when stably transformed with a YAP-based fluorescent construct, the immortalised cells maintained sensitivity to - and were able to report on - the mechanical properties of the surrounding microenvironment. Given the continued interest in MSCs in tissue engineering and regenerative medicine, and the growing appreciation of the roles of mechanical signalling in determining a broad range of cell behaviours, these cells and derivative technologies have potential to benefit scientists wishing to understand the mechanical properties of substrates and scaffolds from a cellular perspective.

## Methods

### Cell culture

Primary human MSCs were harvested from the bone marrow of male and female donors aged 58-80 undergoing knee and hip surgery, under provision of informed written consent. Experiments were performed with relevant National Research Ethics Service and University of Manchester approvals. MSCs were isolated according to established methodology^44^. Primary MSCs were used at passages 3 - 5. Immortalised human MSCs (line Y201) were used as described previously^20^. MSCs were cultured in low glucose (1 g/L) DMEM (Gibco). 293T cells were cultured in high glucose (4.5 g/L) DMEM (Lonza). All media was supplemented with 10% fetal bovine serum (FBS, Labtech) and 1% penicillin/streptomycin cocktail (P/S, Sigma). Investigations into the effects of substrate stiffness were performed on collagen-I coated PA gels (Matrigen).

### Immunofluorescence

Cells were cultured on polyacrylamide hydrogels of defined stiffnesses (Matrigen) at a density of 450 cells/cm^2^ for 3 days. Cells were fixed with 4% paraformaldehyde (PFA, VWR International), permeabilized with Triton X-100 (Sigma), blocked with bovine serum albumin (BSA, Sigma) and stained with DAPI (Sigma), phalloidin (488 and 647; Life Technologies) and primary antibodies, according to manufacturers’ instructions: YAP1 (rabbit, Proteintech); MKL1 (rabbit, Abcam); lamin-A/C (mouse, Santa Cruz Biotechnology); lamin-B1 (rabbit, Proteintech); and GFP (rabbit, Invitrogen). Samples were imaged with secondary antibodies: anti-rabbit 488, 594 and 647; and anti-mouse 594 (all raised in donkey, from Life Technologies). Cells were imaged using an Axio Examiner A1 microscope (Zeiss) with an EC Epiplan-Neofluar 20x/0.5 NA dipping objective lens (Zeiss). Images were processed in ImageJ (version 2.0.0, National Institutes of Health, USA)^45^; CellProfiler (version 2.1.1, Broad Institute, USA)^46^ was used to characterize cell morphometric parameters: cell area; nuclear area; ratios of nuclear to cytoplasmic protein intensities; aspect ratio (ratio of major to minor axes of an ellipse enclosing the cell or nucleus); and circularity (ratio of (4π × area) to square of the perimeter). Images were corrected for background fluorescence by subtracting the mean intensity of a cell-free area from each pixel; all images under comparison in the same experiment had matched exposure and contrast settings.

### Differentiation assays

Osteogenesis was chemically induced by culture in StemXVivo Osteogenic/Adipogenic Base Media (R&D Systems), supplemented with StemXVivo Human Osteogenic Supplement (R&D Systems) and 1% P/S. Adipogenesis was chemically induced in high glucose (4.5 g/L) DMEM with 1 μM dexamethasone, 0.5 mM 3-isobutyl-1-methylxanthine (IBMX), 10 μg/mL insulin and 100 μM indomethacin (all reagents from Sigma), supplemented with 10% FBS and 1% P/S as described previously^47^. MSCs were grown in osteogenic, adipogenic or control media for three weeks on collagen-I coated PA hydrogels for RT-qPCR assays, with seeding densities of 2000 and 1000 cells/cm^2^, for primary and immortalised MSCs respectively (seeding rates were adjusted to account for a greater rate of proliferation in the immortalised cells). For cytochemistry assays, cells were seeded on tissue culture plastic (TCP) at densities of 1200 or 650 cells/cm^2^, for primary and immortalised MSCs respectively.

### Cytochemistry

Adipogenesis was quantified by positive staining of lipid droplets with Oil Red O (Sigma) and osteogenesis by alkaline phosphatase activity (Sigma, following the manufacturer’s protocol). Adipogenic cells were imaged using an EVOS XL Core microscope (Life-Tech) with a Plan PH2 achromatic infinity-corrected 20x lens.

### RT-qPCR

Total RNA was extracted with miRNeasy Mini Kit (Qiagen) and its concentration was measured with a NanoDrop 2000 spectrophotometer (Thermo Scientific). mRNA was reverse transcribed using miScript II RT Kit with the HiFlex buffer (Qiagen) in a Verity Thermal Cycler (Applied Biosystems). Adipogenic (*PPARG* and *CEBPA)* and osteogenic markers (*RUNX2* and *BGLAP)* were quantified with SYBR Select Master Mix (Applied Biosystems) according to the manufacturer’s instructions, with normalisation against *PPIA* (Sigma), in a StepOnePlus Real-Time PCR System (Applied Biosystems). Experiments were performed in technical triplicates and relative gene expression was calculated using the 2-ΔΔCt method^48^. Primers were purchased from PrimerDesign:

*PPARG* (Forward, AACACTAAACCACAAATATACAACAAG; Reverse, GGCATCTCTGTGTCAACCAT)
*CEBPA* (CGG CAACT CTAGTATTTAG GATAAC; CAAATAAAATGACAAGGCACGATT)
*RUNX2* (TTCTCCCCTTTTCCCACTGA; CAAACGCAATCACTATCTATACCAT)
*BGLAP* (CAGCGAGGTAGTGAAGAGACC; TCAGCCAACTCGTCACAGTC)
*PPIA* (ATGCTGGACCCAACACAAA; TTTCACTTTGCCAAACACCA)

### Preparation of reporter constructs

A construct expressing GFP-YAP / RFP-H2B (“GFP-YAP(wt)”) was cloned from a plasmid provided by the Discher laboratory (University of Pennsylvania, USA), with inclusion of peptide T2A to obtain a bicistronic vector (see Supplemental Figure 1). A second construct expressing the same, but with four point mutations to serine residues in YAP (S61A, S109A, S164A and S381G; “GFP-YAP(4SA)”) was generated by site-directed mutagenesis (Agilent QuikChange Lightning kit, used according to the manufacturer’s instructions). Plasmid identities were confirmed by sequencing (GATC Biotech) and analysis with DNADynamo (Blue Tractor Software Ltd.) and BLAST (NCBI, National Institutes of Health, USA). Plasmids were amplified in competent *E. coli* cells, harvested and purified with a HiSpeed Plasmid Maxi Kit (Qiagen), following the manufacturer’s instructions.

### Viral delivery of reporter constructs

Lentiviral envelope plasmid pMD2.G and packaging plasmid psPAX2 were gifts from Didier Trono (Addgene plasmids #12259 and #12260). 293T cells were transiently transfected with the reporter, packaging and envelope plasmids (at a ratio 2:1.5:1) using polyethylenimine (PEI; Merck-Millipore). Virus production was enhanced by addition of 10 mM sodium butyrate (Merck-Millipore) and viral particles concentrated using Vivaspin 20 ultracentrifugation column (Sartorius). Y201 cells were transduced with the viral particles and 8 μg/mL polybrene (Merck-Millipore). Two weeks following infection RFP-positive cells were selected by FACS-sorting.

### Statistical analysis

All data is presented as mean ± SEM. All experiments were performed with four biological primary donors. Data was checked for normal distribution using D’Agostino-Pearson or Shapiro-Wilk tests. Differences in means of normally distributed data were analysed with t-tests (paired when applied to donor-matched samples), otherwise Kruskal-Wallis (KW) or Mann-Whitney (MW) non-parametric tests were used. Analysis was performed using Prism 7 (GraphPad Software Inc.) and Mathematica (Wolfram Research Inc.).

---

**Supplemental Figure 1.**
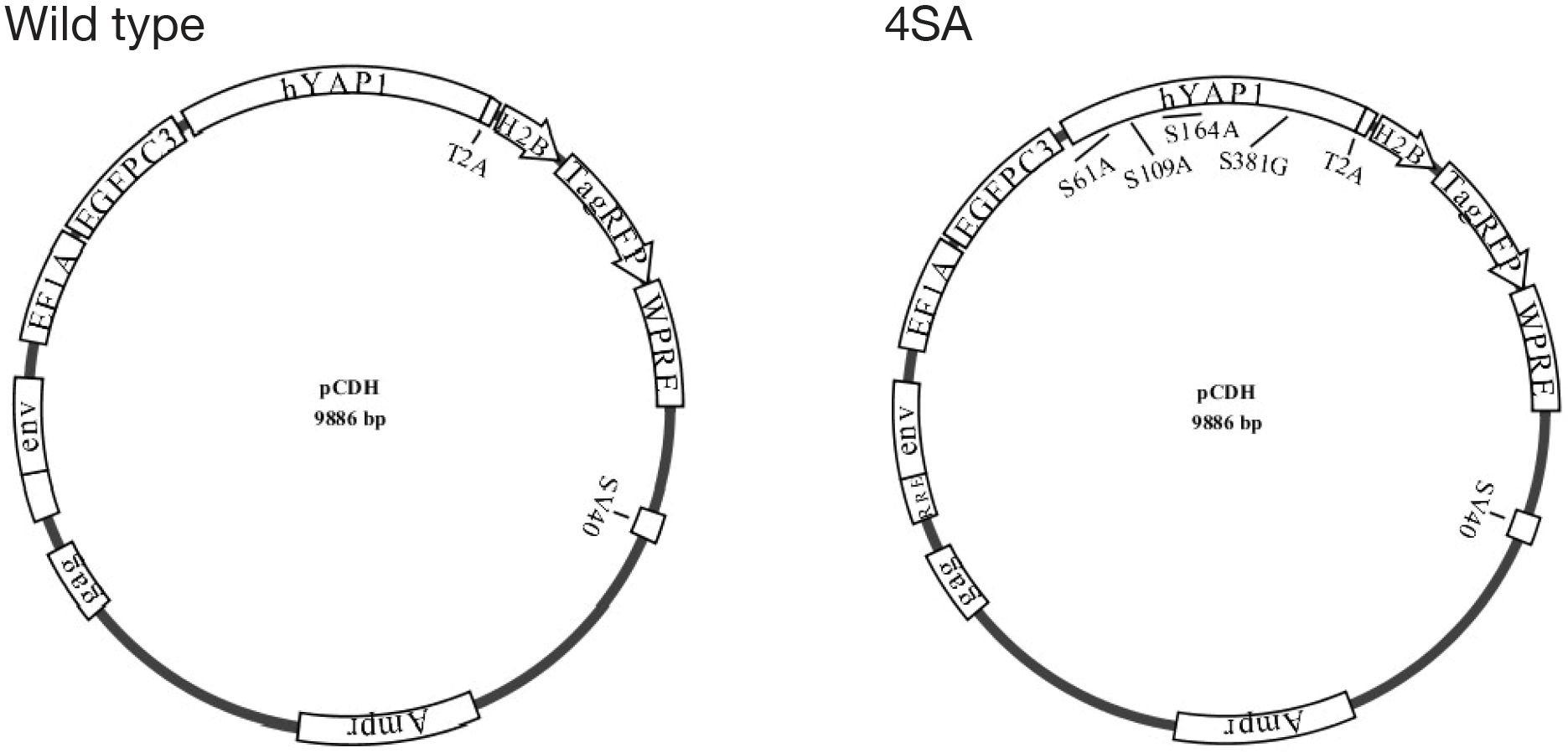
Plasmid maps of constructs used to create reporter cells. Lentiviral backbone plasmids contained the human YAP1 gene tagged to GFP linked with the T2A peptide to histone H2B tagged to RFP. This bicistronic vector allowed for expression of both proteins in the same reading frame, in a stoichiometric manner. Transcription was regulated by an EF1A promoter. The 4SA plasmid has four point mutations on YAP1 serine residues: S61A, S109A, S164A and S381G.

## Acknowledgements

HTJG and JS were funded by a Biotechnology and Biological Sciences Research Council (BBSRC) David Phillips Fellowship (BB/L024551/1). OD was supported by a Wellcome Trust Institutional Strategic Support Fund (097820/Z/11/B). PG was funded by the National Centre for the Replacement, Refinement and Reduction of Animals in Research (NC3Rs) and Arthritis Research UK (ARUK). Microscopy was carried out at the Wellcome Trust Centre for Cell-Matrix Research (WTCCMR; 203128/Z/16/Z) Imaging Core Facility. The authors thank: Tim Board (Wrightington Hospital) for provision of tissue samples for MSC isolation; Mike Jackson and Gareth Howell (University of Manchester Flow Cytometry Core Facility); and Robert Pedley for assistance with viral protocols.

## Author Contributions

Investigation, AGT, JES, AW, OD, HTJG; Formal Analysis, AGT, JES and JS; Writing - Original Draft, AGT; Writing - Review & Editing, AGT, JES, AW, OD, HTJG, PG, KB, SMR and JS; Visualization, Project Administration and Funding Acquisition, PG, KB, SMR and JS.

## Additional Information

*Competing financial interests: The authors declare no competing financial interests*.

